# Convergent evolution of parrotfish beaks enables stress redistribution during high-force feeding

**DOI:** 10.64898/2026.04.27.721161

**Authors:** Ritika Raj Menghani, Sean Anthony Trainor, Kory M. Evans, Raudel Avila

## Abstract

Convergent evolution is often interpreted as evidence that similar ecological challenges select for optimal functional solutions, yet the biomechanical mechanisms underlying such convergence remain poorly resolved. Parrotfishes independently evolved fused, beak-like dentition multiple times, a trait associated with feeding on hard substrates such as coral, but its advantage has not been quantified. Here, we combine finite element analysis with shockwave modeling to evaluate the performance of the beak during feeding. Across five species spanning independent evolutionary origins of beaked and non-beaked dentitions, we find that jaw opening is governed primarily by geometric scaling, with minimal influence from dentition morphology. In contrast, under stabilized biting, fused dentition reduces the rate of stress accumulation relative to discrete teeth, indicating a functional advantage under constrained loading. Shockwave modeling further shows that tooth geometry and stacking regulate stress propagation: smooth profiles reduce stress concentration, while stacked architectures localize stresses within the tooth material and limit transmission into the surrounding bone. These results suggest that convergent beak evolution in parrotfish reflects repeated optimization for stress management under constrained and impact loading, and reveals a general principle by which biological structures control internal force transmission during high-force feeding rather than maximize strength alone.

## 1 INTRODUCTION

Convergent evolution is a striking pattern that occurs when distantly related species independently evolve traits and appearances that closely resemble one another. Convergence has manifested in the elongated snouts of African and South American electric fishes^1,2^, the crab-like body shapes of several crustacean lineages^3,4^, and the succulence of plant appearances in dry arid environments^5^ with countless more examples. The driving forces behind convergent evolution are diverse but are thought to frequently indicate similarities in function or ecology. As a result, convergent evolution has been described as some of the strongest evidence for the effect of natural selection on shaping organismal phenotypes.

Among ray-finned fishes (Actinopterygii), diet has been shown to act as a powerful driving force for convergent evolution and has produced independently evolved trait similarities across many trophic guilds. Many of these traits are closely associated with the morphology of the skull and head including tooth reduction and jaw adductor muscle reductions in planktivorous surgeonfishes^6^ and in the myriad of convergent head shapes in the Lake Tanganyika and Lake Malawi cichlids^7^. Among the ray-finned fishes, parrotfishes (Scarini) have recently been found to exhibit striking patterns of convergent evolution in the presence of their beak-like dentition where hundreds of small teeth have been coalesced into a complex series of overlapping tooth rows that together form robust tooth plates that allow these species to scrape and excavate hard surfaces like rocks and coral skeletons. Using their robust beaks, parrotfishes function as ecosystem engineers and perform important roles in grazing, bioerosion and dispersal of coral algal symbionts and are thus considered essential for maintaining the health of coral reef ecosystems.

A study by Evans, et al.^8^ found that beaked dentition arose once in the “reef” clade of parrotfishes that includes *Scarus, Chlorurus, Hipposcarus, Cetoscarus*, and *Bolbometapon* and again in the “seagrass” clade that includes the Atlantic Ocean endemic *Sparisoma*. While the presence of coalesced teeth in parrotfishes represents a striking pattern of convergence, there is also substantial diversity in the beak shapes distributed across the different parrotfish species that have been hypothesized to correspond to differences in foraging ecology. When their foraging habitats were initially described, they were assumed to be indiscriminate generalist herbivores of the coral reef that employed a wide range of foraging behaviors to graze. However, recent work has shown that parrotfishes exhibit immense dietary specificity and specialization in not only what they feed on, but also how they feed^9,10^. Studies have shown that parrotfishes typically employ three stereotyped foraging behaviors during feeding. 1) Browsing: species use jaws to remove macroalgae or seagrass without scraping or scarring the substrate. 2) Excavating: species use robust jaws to remove pieces of substrate during feeding. 3) Scraping: species feed on the surface of the substrate with little to no scarring. Browsing has been hypothesized to be the ancestral foraging mode of all parrotfishes while excavation and scraping are more derived^11^. These different foraging groups are typified for several cranial adaptations, in scrapers, many species exhibit an intra-mandibular joint in addition to their typical quadrato-articular joint that allows them to gain additional degrees of flexibility in their lower jaws which may allow scraping species to feed significantly faster than excavating species, sometimes taking twice as many bites on substrate within a given observation timeframe^12–14^. The beaks of scrapers are often more gracile in appearance than the excavators and exhibit a diverse range of shapes. Some excavating species lack the additional jaw joint that scrapers have and instead possess robust beaks that are often fully cemented over which they use to take strong deep bites out of substrate^8,11^. While most parrotfish species feed on dead corals, some excavating species are known to feed on living corals and have been documented removing large amounts of reef biomass during feeding bouts^9,10,13,15–17^. While the jaws of scraping parrotfishes are diverse in shape, there are strong patterns of convergence in the jaws of excavating species suggesting that there may be stronger selection towards a shared phenotype or fewer functional solutions for building excavating jaws^8^. Browsing is believed to be the ancestral foraging state of all parrotfishes^8,11,18,19^. Browsing species typically lack beaks and instead possess conical teeth with various levels of coalescence. These species do not feed on hard substrates and instead mostly target aquatic vegetation and macroalgae. Despite decades of ecological and morphological study, it remains unclear whether these differences in beak shape and coalescence translate into meaningful differences in biomechanical performance.

Finite element analysis (FEA) provides a quantitative framework for evaluating the mechanical performance of complex biological structures under controlled loading conditions. Previous studies have used FEA to investigate functional consequences of morphological variation, including patterns of convergent evolution in three-dimensional structures such as skulls and jaws^20,21^. In parrotfishes, fused beak-like dentition has been widely hypothesized as an adaptation for durophagy; however, its biomechanical role remains unresolved. In particular, it is not known whether fused dentition provides a measurable advantage during loading conditions associated with feeding on hard structures. Here, FEA is used to quantify stress distributions in parrotfish jaws and to test for functional differences between beaked and non-beaked morphologies. The analysis includes species spanning two independently evolved beak lineages: *Scarus forsteni, Sparisoma viride* and *Cetoscarus ocellatus*, which exhibit fused dentition, and *Cryptotomus roseus* and *Calotomus japonicus*, which retain discrete teeth (**Fig. 1a**). In addition, simplified reduced-order models are used to examine the mechanical roles of tooth geometry and stacking under transient shockwave loading. This approach enables the first direct evaluation of whether convergent beak morphology reflects a biomechanical advantage associated with stress redistribution and impact loading.

**Figure 1.**
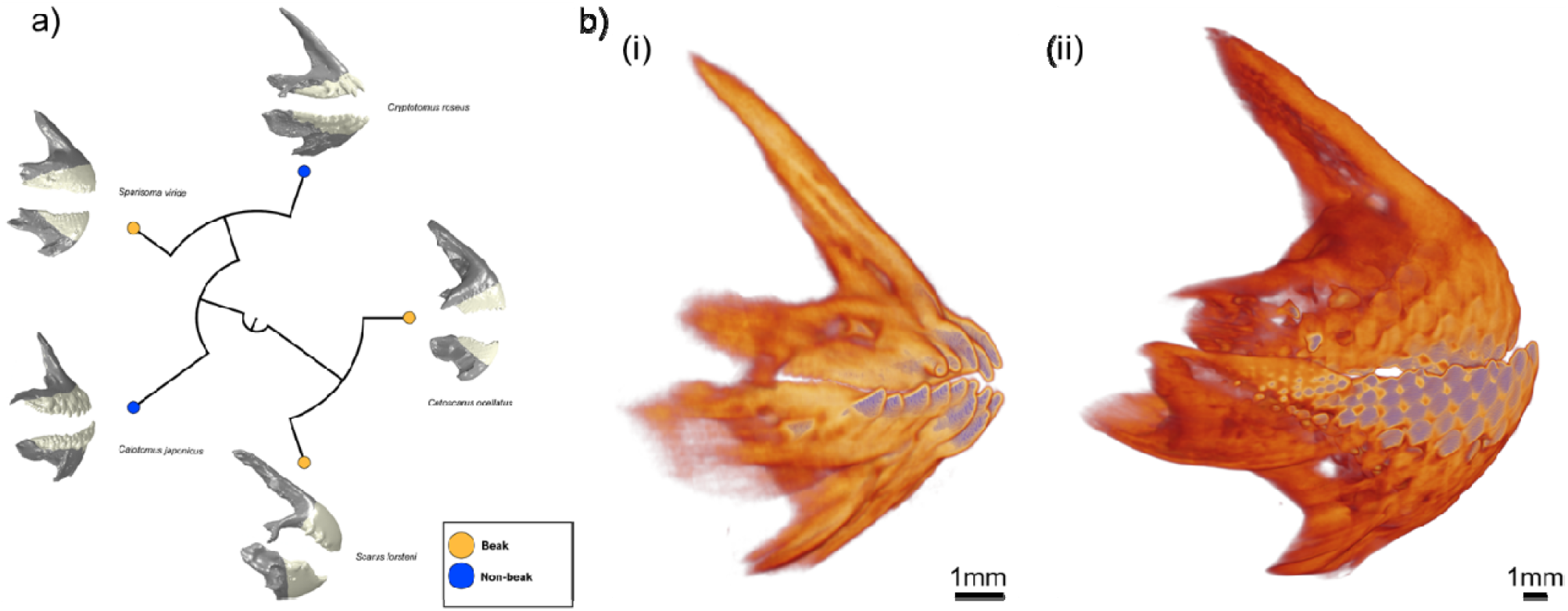
Convergent evolution of the beak structure in parrotfishes. a) The evolutionary relationships of five parrotfish species (*Cryptotomus roseus, Cetoscarus ocellatus, Scarus forsteni, Sparisoma viride, and Calotomus japonicus*) are shown. The grey regions represent bone, and the white regions represent teeth. Beak-forming species (yellow nodes) and non-beak species (blue nodes) are distributed across the phylogeny, demonstrating that the beak structure evolved independently across species. b) Micro-CT Imaging for (i) *Cryptotomus roseus* showing non beak morphology (ii) *Sparisoma viride* showing the beak like geometry.

## 2 MATERIALS AND METHODS

### 2.1 Digital Reconstructions

Five parrotfish species were selected to span independent evolutionary origins of beak-like dentition and a range of dental morphologies: *Cryptotomus roseus, Scarus forsteni, Calotomus japonicus, Sparisoma viride* and *Cetoscarus ocellatus*. This selection enabled controlled comparison of mechanical response across independently derived morphologies. Specimens were imaged using micro-computed tomography (micro-CT) to obtain high-resolution anatomical geometries, with voxel sizes on the order of tens of micrometres. Specimens were scanned using a Bruker Skyscan 1273 and segmented in Amira^22^ to isolate the left premaxilla and dentary bones of the upper and lower jaws of each specimen. Jaw bones were then additionally segmented to isolate the tooth plate and the surrounding jaw bone (**Figure 1b**). Segmented structures were then exported as three-dimensional mesh models for downstream analyses. Resulting geometries were processed to generate closed surfaces suitable for finite element meshing. Geometric processing was limited to removal of imaging artefacts and closure of small voids, without altering overall morphology (**Supplementary Fig. 1**).

### 2.2 Finite Element Analysis – Jaw Opening and Jaw Closing

The segmented CT geometries were processed to remove surface intersections and close small voids, yielding suitable three-dimensional volumes suitable for meshing. COMSOL Multiphysics generated second-order tetrahedral meshes (C3D10), with element counts ranging from 75,000 to 140,000 depending on dentition complexity. The models were then exported to ABAQUS for mechanical analysis (**Supplementary Fig. 1**). Table 1 reports the mass moment of inertia, mass, volume, and element count for each model.

**Table 1.** Mass, volume and element count for all models.

| Parrotfish | Region | Mass Moment of Inertia (kg-mm<sup>2</sup>) | Mass (kg) | Volume (mm<sup>3</sup>) | Number of Elements (C3D10) |
| --- | --- | --- | --- | --- | --- |
| *Cryptotomus roseus* | Premaxilla | 9.64E-05 | 2.03E-05 | 8.92 | 92,700 |
|  | Dentary | 8.65E-05 | 2.55E-05 | 10.8 | 138,577 |
| *Calotomus japonicus* | Premaxilla | 7.78E-03 | 4.75E-04 | 191 | 83,875 |
|  | Dentary | 7.04E-03 | 4.61E-04 | 193 | 77,646 |
| *Scarus forsteni* | Premaxilla | 7.18E-03 | 3.48E-04 | 136 | 86,671 |
|  | Dentary | 3.95E-03 | 2.67E-04 | 112 | 79,047 |
| *Sparisoma viride* | Premaxilla | 8.48E-03 | 5.84E-04 | 242 | 113,145 |
|  | Dentary | 1.43E-02 | 6.64E-04 | 287 | 122,905 |
| *Cetoscarus ocellatus* | Premaxilla | 1.10E-01 | 1.88E-03 | 843 | 86,889 |
|  | Dentary | 9.94E-02 | 2.49E-03 | 1,060 | 90,419 |

Linear elastic material properties were assigned to bone and tooth regions (Table 2). Due to limited experimental data for parrotfish hard tissues, identical material properties were applied across all species to enable direct comparison of mechanical response. Because the study focuses on relative differences between morphologies, the use of uniform material properties does not affect the comparative interpretation of stress distributions. Bone properties were based on cortical human bone, while tooth density was approximated using fluorapatite, consistent with the mineral composition of enameloid^23^. Two loading conditions were analyzed using static finite element simulations: (i) jaw opening, in which the structure deforms under applied load, and (ii) jaw closing, in which displacement is constrained to evaluate stress development under stabilized biting.

**Table 2.** Material properties for bone and tooth regions.

| Region | Young's Modulus (GPa)<br>(E) | Poisson's Ratio<br>(ν) | Density (kg/m<sup>3</sup>)<br>(ρ) | Reference |
| --- | --- | --- | --- | --- |
| Bone | 20 | 0.3 | 1850 | 24–26 |
| Teeth | 110 | 0.27 | 3180 | 23,27,28 |

### 2.3 Finite Element Analysis – Impact and Wave Propagation

Biting into hard substrates generates transient stress waves that propagate through the parrotfish jaw. Understanding how parrotfishes sustain repeated impact loading provides insight into the mechanical role of tooth geometry and arrangement. To isolate these effects, a simplified model representing a section of the jaw was constructed as a cuboid with distinct bone and tooth regions. Two configurations were analyzed: one to evaluate the influence of tooth shape on stress generation, and another to examine stress wave propagation in stacked tooth architectures. All simulations employed an explicit dynamic formulation to resolve transient wave behavior.

Fish teeth exhibit substantial geometric variation^29–32^; therefore, a simplified model was constructed to evaluate the effect of tooth shape on transient stress propagation during biting. The model includes cuboid, frustum, spade, capped pyramid and capped sphere geometries, along with a toothless control (**Supplementary Fig. 2a(i–v)**). Each configuration consists of a cuboid base 12 mm in length with a cross-section of 2 mm × 2.5 mm (**Supplementary Fig. 2b**), with dimensions approximated from the dentary of *Sparisoma viride*. Hexahedral elements (C3D8) were used for all geometries except the spade profile, which required second-order tetrahedral elements (C3D10) to capture geometric complexity.

For the second study, a stacked tooth architecture was constructed using the spade profile, selected for its similarity to parrotfish tooth geometry^8,11,33^. The geometry was generated by sequentially offsetting the profile to form a continuous tooth region comprising 32 teeth. The model consists of a cuboid base with a 1 mm × 1 mm cross-section and a total length of 8.2 mm (**Supplementary Fig. 2c**), with a tooth volume fraction of 2.4%. The mesh employs second-order tetrahedral elements (C3D10). Differences in wave propagation between tooth and bone regions arise from their distinct material properties. The longitudinal wave speed (c) in an elastic medium was estimated using

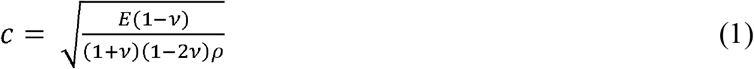

where (E) is the elastic modulus, (*ν*) is Poisson’s ratio and (*ρ*) is the density. Using the material properties defined in Table 2, wave speeds were estimated to be higher in tooth material (approx. 6,574 m s□^1^) than in bone (approx. 3,814 m s□^1^).

## 3 RESULTS

### 3.1 Jaw Opening

During feeding, bite forces applied at the teeth induce opening of the jaw. To enable direct comparison across species, identical loading conditions were applied to all models. A 48 N load was distributed across the tooth tips in the Y direction, while the posterior region was fully constrained to represent a hinge-like boundary condition (**Fig. 2a(i)**). The 48N value was chosen because it fits within the range of estimated bite force for some parrotfish species^34^. Lateral motion was restricted by fixing displacement in the Z direction to enforce symmetry. **Figure 2a(ii–vi)** shows the resulting displacement contours, with deformed geometries overlaid on the undeformed configuration.

**Figure 2.**
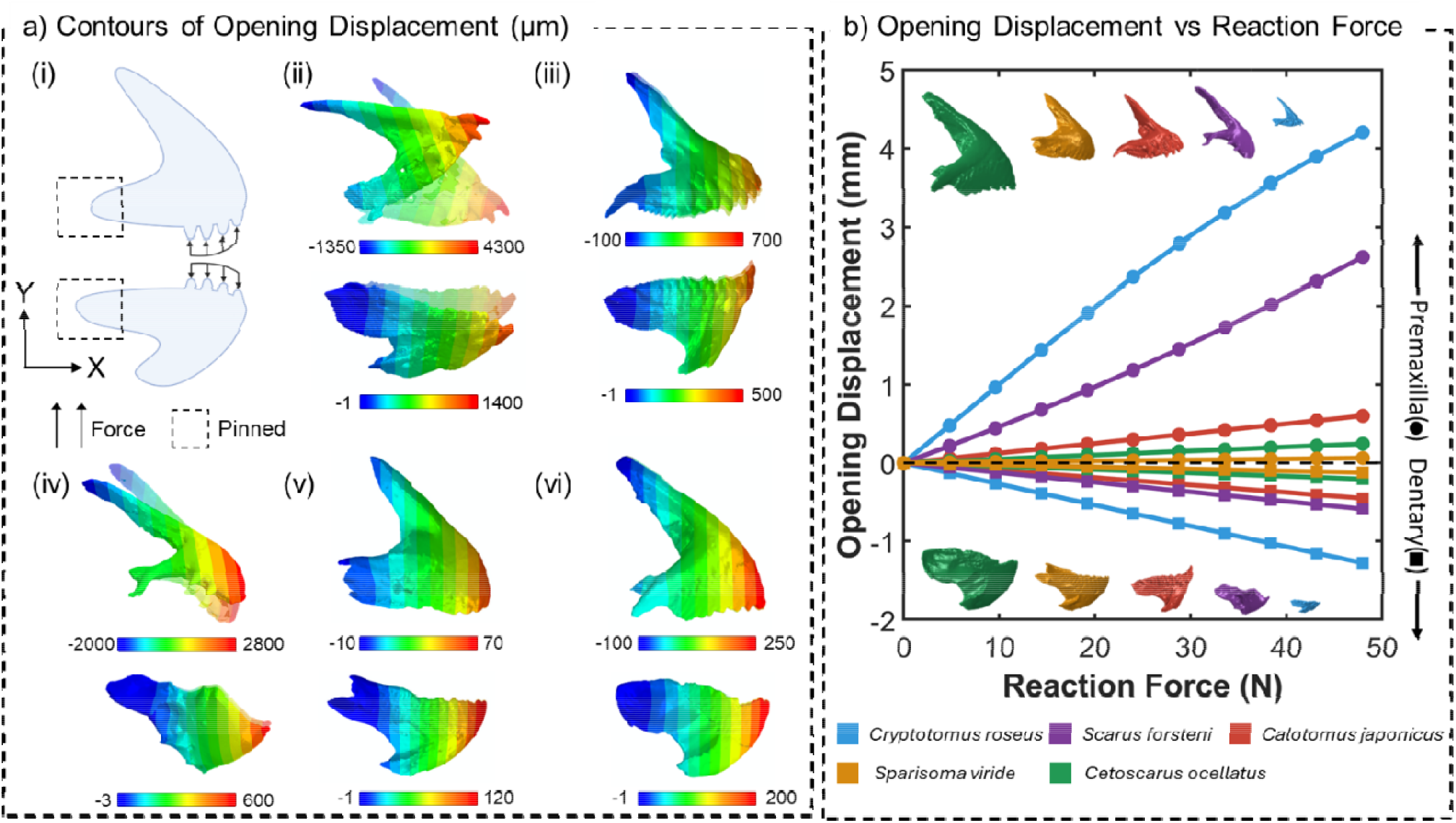
Jaw opening simulations. (a) Boundary conditions and displacement contours. (i) A 48 N load was applied at the tooth region while the posterior end was constrained to represent a hinge-like condition (Created i https://BioRender.com). Displacement fields are shown for (ii) *Cryptotomus roseus*, (iii) *Scarus forsteni*, (iv) *Calotomus japonicus*, (v) *Sparisoma viride* and (vi) *Cetoscarus ocellatus*, with deformed geometries overlaid on the undeformed configuration. *Cryptotomus roseus* exhibits the largest opening displacement. (b) Tip openin displacement as a function of reaction force for all models. Geometries are shown to scale. Displacement increases with decreasing model size, with reduced displacement observed in *Sparisoma viride* due to differences in posterior geometry.

The force–displacement response indicates that opening deformation is governed primarily by geometric scaling rather than dentition morphology. **Figure 2b** plots tip displacement as a function of reaction force for each model, with geometries shown to scale. In all cases, the reaction force reaches 48 N, consistent with the applied load. Displacement scales primarily with overall geometry: *Cryptotomus roseus* exhibits the largest displacement (approx. 4 mm), followed by *Scarus forsteni* (approx. 2.5 mm), despite differences in dental morphology. The remaining species exhibit displacements below 1 mm. An exception occurs between *Sparisoma viride* and *Cetoscarus ocellatus*, where *Sparisoma viride* exhibits smaller displacements (0.06– 0.12 mm) despite comparable or smaller size. This difference is attributed to variation in posterior geometry, where a proportionally larger hinge region increases resistance to rotation.

To quantify this behavior, Table 1 reports the mass moment of inertia about the Z axis for each model. The values span multiple orders of magnitude, with *Cryptotomus roseus* exhibiting the lowest inertia and *Sparisoma viride* and *Cetoscarus ocellatus* the highest. This trend aligns with the displacement results and indicates that resistance to rotation, rather than dentition morphology, governs opening behavior. Table 3 further reports effective stiffness and strain energy. Effective stiffness, defined as the ratio of reaction force to tip displacement, increases with model size, while strain energy, obtained from the force–displacement response, decreases. Together, these results show that global deformation during jaw opening is controlled by geometric scaling, with minimal influence from beak morphology.

**Table 3.** Effective stiffness and strain energy for all models under jaw opening.

| Parrotfish | Region | Effective Stiffness ($\times 10^8$ N/mm) | Strain Energy (N-mm) |
| --- | --- | --- | --- |
| *Cryptotomus roseus* | Premaxilla | 0.1142 | 108.9830 |
|  | Dentary | 0.3754 | 30.9096 |
| *Calotomus japonicus* | Premaxilla | 0.7984 | 14.4111 |
|  | Dentary | 1.0602 | 10.8123 |
| *Scarus forsteni* | Premaxilla | 0.1830 | 58.9995 |
|  | Dentary | 0.8137 | 14.1994 |
| *Sparisoma viride* | Premaxilla | 7.6524 | 1.5048 |
|  | Dentary | 3.9134 | 2.9408 |
| *Cetoscarus ocellatus* | Premaxilla | 1.9590 | 5.8694 |
|  | Dentary | 2.3410 | 4.9175 |

The large displacement observed for *Cryptotomus roseus* requires careful interpretation. The predicted opening displacement (approx. 4 mm) is comparable to the characteristic dimensions of the model (approx. 7.5 mm; **Supplementary Fig. 3**), indicating that the applied load approaches the limits of the linear deformation regime assumed in the analysis. This behavior is consistent with the substantially lower mass moment of inertia for *Cryptotomus roseus*, which reduces resistance to rotation under the prescribed loading conditions. The force–displacement response for this model exhibits deviation from linearity beyond approximately 24 N, further suggesting that geometric effects become significant at higher loads. Because identical loading conditions are applied across all species to enable direct comparison, the resulting response for smaller geometries reflects a scaling limitation rather than a physiological prediction. Accordingly, the results for *Cryptotomus roseus* at higher loads are interpreted qualitatively. Future studies should incorporate species-specific loading conditions to identify the range of linear response and the onset of nonlinear behavior under physiologically relevant opening forces. Taken together, jaw opening is governed primarily by geometric scaling, including model size, hinge region geometry and rotational inertia, with minimal influence from dentition morphology.

### 3.2 Jaw Closing

In addition to jaw opening, we also modeled constrained jaw closing in the premaxilla and dentary. To simulate this condition, a 48 N load was applied to the tooth tips in the Y direction, while the superior surface of the premaxilla (or inferior surface for the dentary) was fully constrained to prevent vertical displacement (**Fig. 3a(i)**). Lateral motion was restricted by fixing displacement in the Z direction. Under these boundary conditions, deformation is minimized and the response is characterized by stress development.

**Figure 3.**
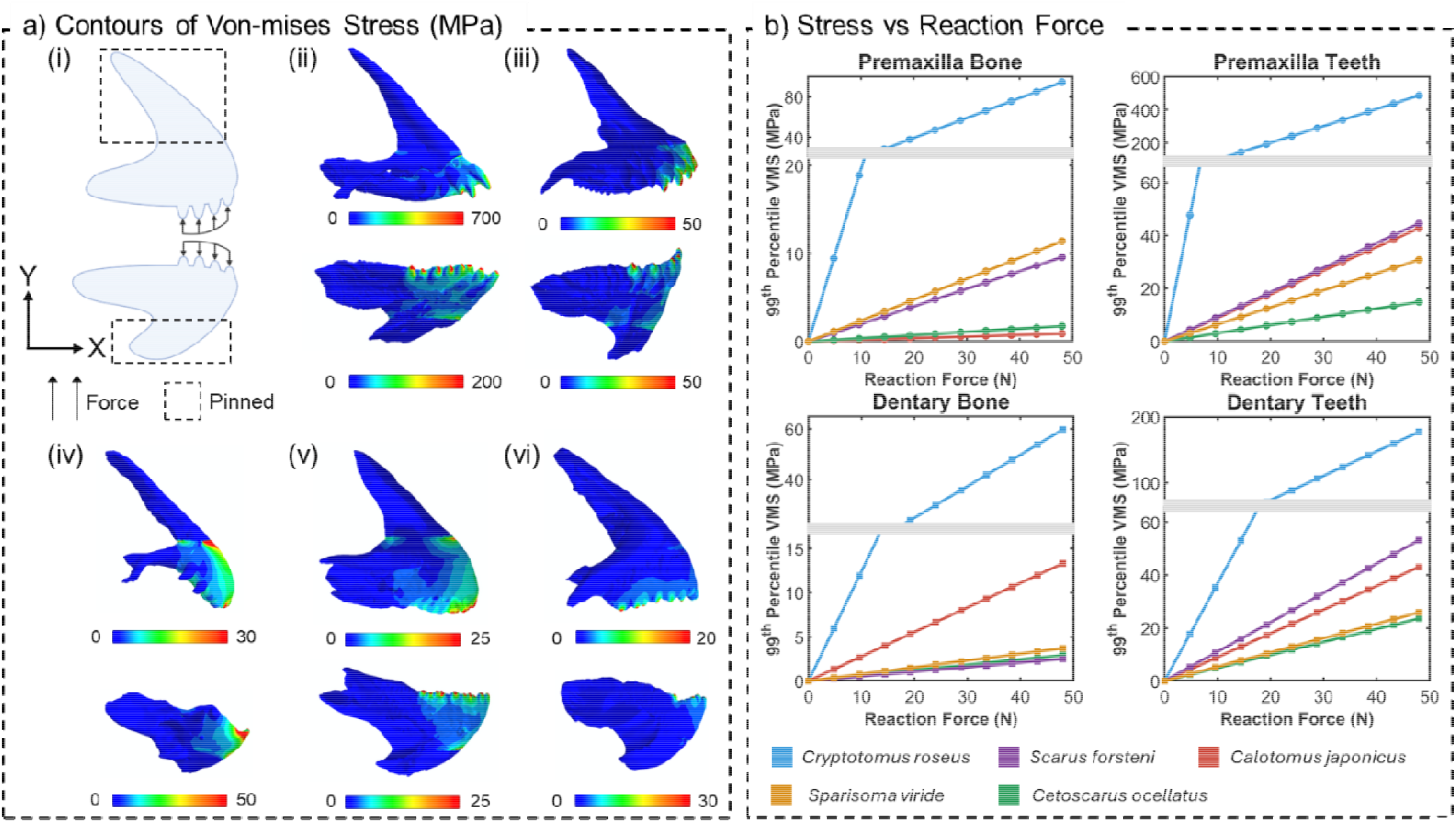
Jaw closing simulations. (a) Boundary conditions and von Mises stress contours. (i) A 48 N load was applied at the tooth region while the superior surface was constrained to prevent vertical displacement (Created in https://BioRender.com). Stress distributions are shown for (ii) *Cryptotomus roseus*, (iii) *Scarus forsteni*, (iv) *Calotomus japonicus*, (v) *Sparisoma viride* and (vi) *Cetoscarus ocellatus*. Elevated stresses localize primarily within the tooth region. (b) 99th-percentile von Mises stress as a function of reaction force for (i) premaxilla tooth, (ii) dentary tooth, (iii) premaxilla bone and (iv) dentary bone regions. A break in the y-axis highlights trends amon lower-stress species. *Cryptotomus roseus* exhibits the highest stress levels, whereas *Cetoscarus ocellatus* maintains consistently lower stresses.

**Figure 3a(ii–vi)** shows the resulting von Mises stress distributions, with elevated stresses localized primarily within the tooth region across all species. A consistent trend emerges between dentition morphologies: non-beak species exhibit higher stresses than beak-forming species. *Cryptotomus roseus* shows the highest stress levels (approx. 200–700 MPa), followed by *Calotomus japonicus* (approx. 50 MPa). In contrast, beak species generally maintain lower stresses, with *Sparisoma viride* and *Cetoscarus ocellatus* remaining below 50 MPa. *Scarus forsteni* exhibits intermediate stress levels (approx. 50 MPa), likely reflecting its smaller geometry. These results indicate that fused dentition reduces stress accumulation under constrained loading conditions.

To quantify stress development while limiting sensitivity to localized numerical artifacts, the 99th-percentile von Mises stress was used as a representative metric. **Figure 3b** plots this quantity as a function of reaction force for each model, and Table 4 reports the corresponding slopes (MPa/N). Across all species, stress increases more rapidly in the tooth region than in the bone, as indicated by consistently higher stress–force slopes. *Cryptotomus roseus* exhibits the highest stress levels, reaching approximately 0.5 GPa. Because these values exceed those of the remaining species by more than an order of magnitude, a break in the Y axis is introduced to visually resolve trends in the lower-stress cases. *Calotomus japonicus*, the second non-beak species, also exhibits elevated stresses in the tooth region (approx. 0.04 GPa). In contrast, beak-forming species maintain lower stress levels, with *Cetoscarus ocellatus* remaining below approximately 0.025 GPa across the loading range. The stress–force slopes in the tooth region follow trends consistent with geometric scaling, with higher values observed in smaller models and lower values in larger models. *Calotomus japonicus* and *Scarus forsteni* exhibit comparable slopes, consistent with their similar sizes. In contrast, slopes in the bone region remain low across all species and show no clear dependence on size or dentition morphology. This behavior indicates that stresses are preferentially localized within the tooth material during jaw closing, while the underlying bone experiences comparatively limited loading. Taken together, these results show that stress accumulation during constrained biting is governed by both geometry and dentition morphology, with fused dentition reducing the rate of stress development and promoting localization within the tooth region.

**Table 4.**
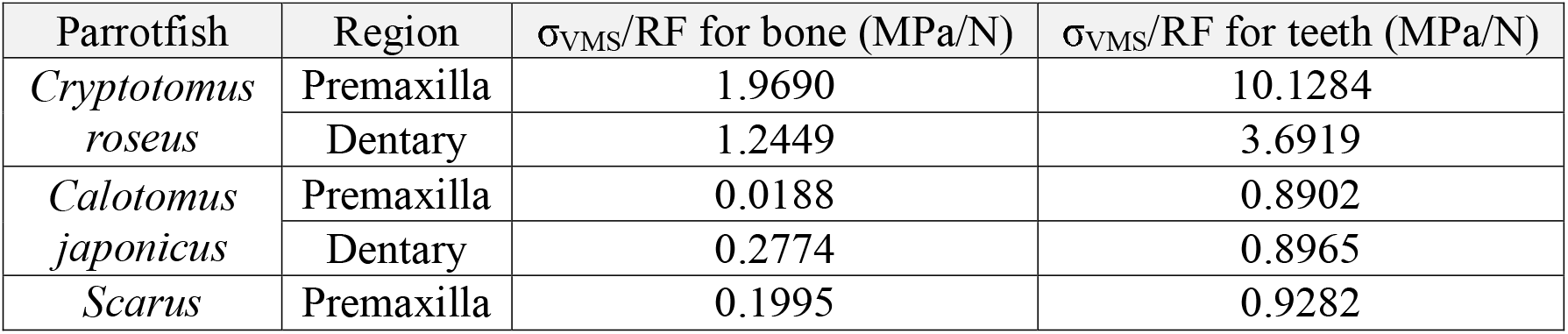
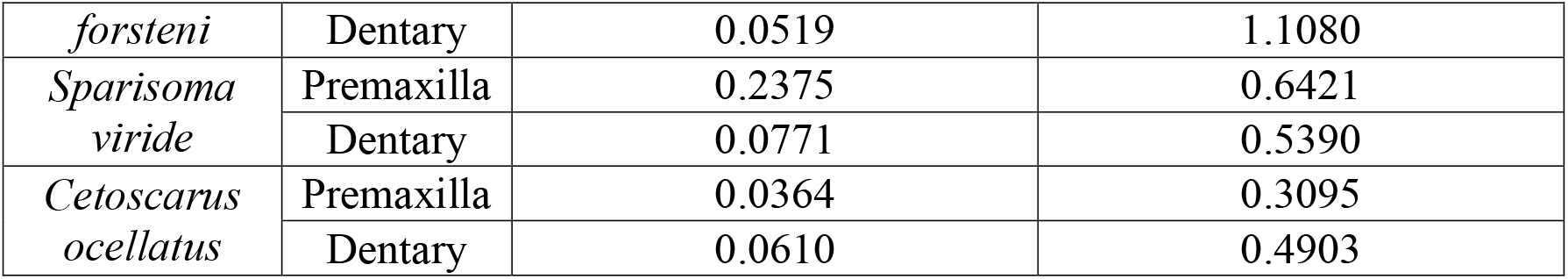
Slope of 99th-percentile von Mises stress versus reaction force for bone and tooth regions of the premaxilla and dentary.

### 3.3 Tooth geometry and stress wave propagation

To evaluate the effect of tooth geometry on transient stress propagation, a reduced-order model was constructed using a representative cross-section of the jaw with a single tooth profile (cuboid, frustum, capped pyramid, capped sphere, and spade) positioned at one end (**Fig. 4a**). A straight cuboid base was used to isolate geometric effects and enable direct comparison of wave propagation across shapes. An identical impulse was applied to the tooth surface in each case (**Fig. 4b**), scaled to maintain a constant total force of 0.5 N across all models. This impulse profile was selected to ensure a shockwave would propagate based on the geometry and material of the model. The tooth was positioned near the edge of the base to approximate loading at the jaw boundary, with a material transition from tooth to bone located at mid-length (**Fig. 4c**). Lateral motion was constrained and the base was fixed to promote one-dimensional (longitudinal) propagation along the length.

**Figure 4.**
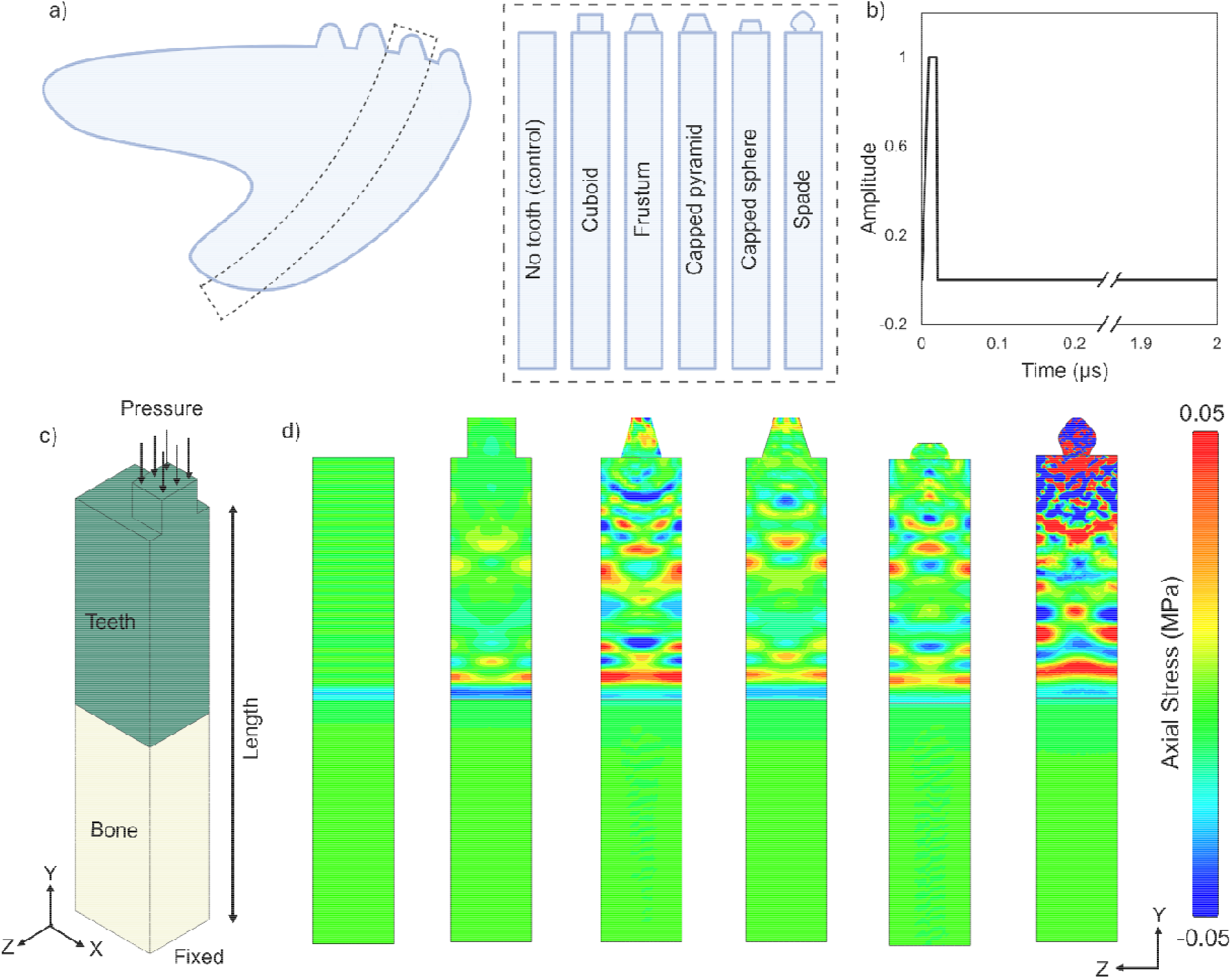
Finite element simulations of tooth impact across geometric profiles. (a) Simplified model geometry derived from a representative section of a parrotfish dentary, used to compare five tooth profiles (cuboid, frustum, capped pyramid, capped sphere and spade) and a flat, no-tooth control. (Created in https://BioRender.com) (b) Temporal profile of the applied impulse load used to simulate biting. (c) 3D Model geometry and boundar conditions. The base was fixed, lateral faces were constrained with roller conditions, and a pressure waveform was applied at the tooth surface. The pressure magnitude was adjusted across cases to maintain a constant total force. (d) Axial stress contours at mid-length wave propagation. No tooth control produces a planar wavefront, whereas all tooth geometries introduce spatial heterogeneity. The spade geometry generates the largest tensile and compressive regions due to reduced contact area and increased applied pressure.

Figure 4d shows axial stress contours at the time corresponding to mid-length wave propagation. Arrival times are similar across tooth geometries (approx. 0.9 μs), with a slightly earlier arrival in the toothless control (approx. 0.8 μs) due to reduced propagation distance. The control case produces a planar wavefront, whereas all tooth geometries introduce spatial heterogeneity, generating localized tensile–compressive regions. The magnitude of this heterogeneity varies with geometry, with sharper or more tapered profiles producing greater stress asymmetry.

To isolate geometric effects from differences in applied pressure, stresses were normalized by the input pressure (**Fig. 5**). The normalized stress profiles show that the cuboid geometry produces the highest stress amplitudes, whereas the spade geometry produces the lowest. The capped sphere and pyramid geometries exhibit responses comparable to the frustum and are shown in **Supplementary Fig. 4**. These results indicate that smooth and tapered geometries reduce stress concentration and disrupt coherent wave transmission relative to sharp-edged profiles.

**Figure 5:**
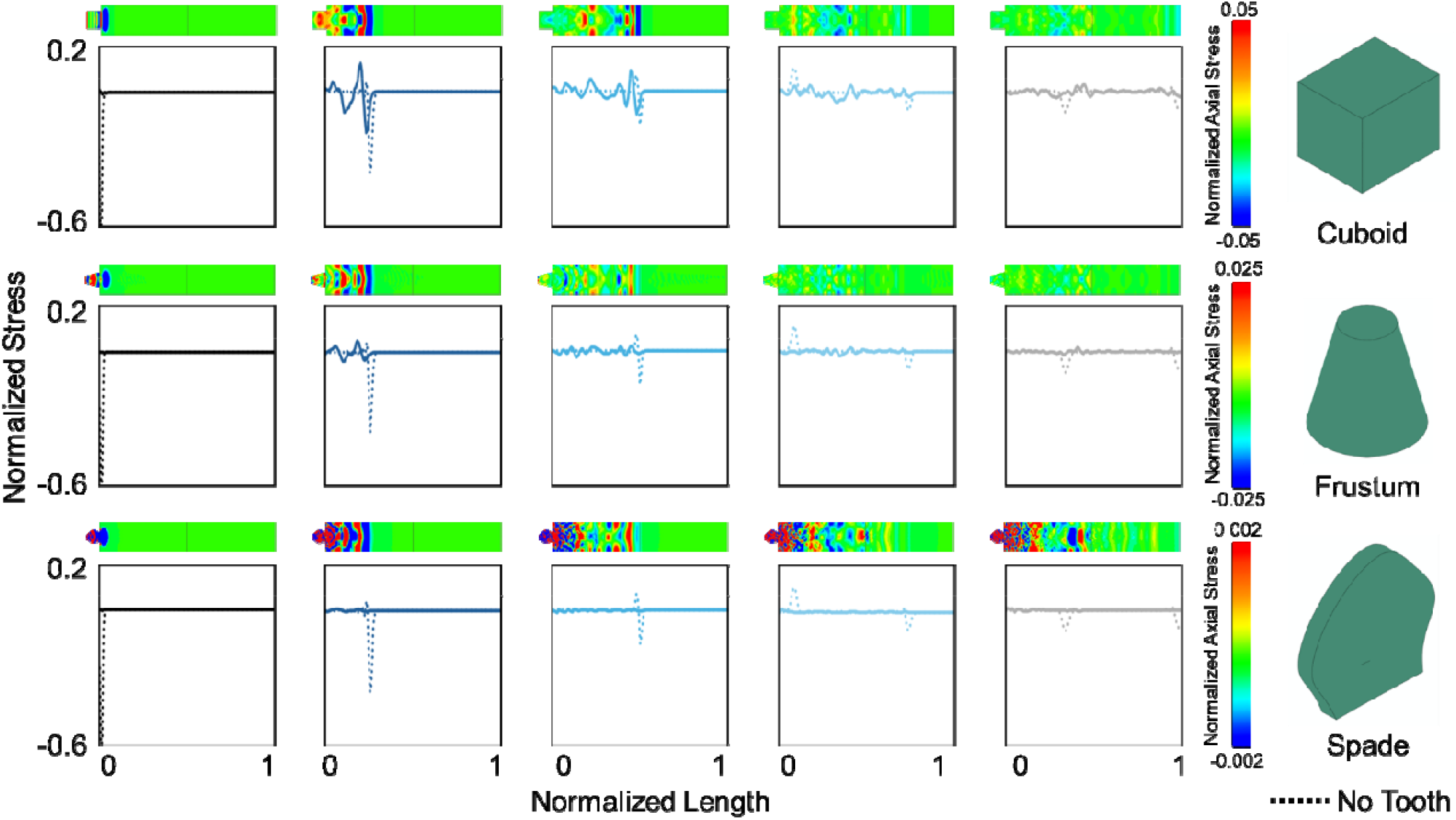
Normalized axial stress distributions for different tooth geometries. Axial stress along the model length is normalized by the applied pressure to enable comparison across geometries. Solid lines show stress profiles for each tooth geometry (cuboid, frustum and spade), with the No tooth control shown as a dashed line. Columns correspond to tooth geometry and rows to successive time points; normalized axial stress contours are shown above each profile. The cuboid geometry produces the highest normalized stress amplitudes, whereas the spade geometr produces the lowest. Sharper geometries generate higher stress concentrations, while smoother, curved profiles reduce stress magnitude and disrupt coherent wave propagation. The spade geometry exhibits increased spatial variation with alternating tensile–compressive regions, consistent with enhanced wave scattering.

At the tooth–bone interface, a mismatch in material properties produces partial reflection and attenuation of the propagating wave. Stress amplitudes decrease by approximately 94% for the cuboid, 86% for the frustum and 55% for the spade geometry. In the toothless control case, a pronounced reflected tensile wave forms at the interface, reducing the forward compressive wave amplitude by approximately 61%. These results show that tooth geometry and material discontinuities jointly control both stress magnitude and wave structure during impact loading.

### 3.4 Tooth stacking

Parrotfishes exhibit a stacked tooth architecture in which successive teeth form a continuous pathway through the jaw (**Fig. 6a(i)**). To evaluate the mechanical role of this arrangement under impact loading, the stacked model (Section 2.3) was subjected to a pressure impulse of 3.5 MPa representative of biting^35^ (**Fig. 6a(ii)**). Three material configurations were analyzed: all bone, all tooth and a composite tooth–bone structure. The all-bone and all-tooth cases define limiting behaviors, while the composite case represents the biological parrotfish configuration. A pressure impulse was applied at the top surface, the base was fixed, and lateral motion was constrained to promote one-dimensional propagation (**Fig. 6b**). Simulations were performed over 8 μs to capture multiple wave reflections. The all-tooth configuration exhibits the fastest wave arrival (approx. 1.28 μs) and the highest stress amplitudes, reflecting the higher stiffness of tooth material. In contrast, the all-bone and composite cases show slower wave propagation (approx. 2.2 μs and 2.1 μs, respectively) and reduced peak stresses. The all-tooth case produces a peak compressive stress of approximately 2.15 MPa, whereas the all-bone and composite cases remain near 1–1.3 MPa. In the composite configuration, the mismatch in material properties at the tooth–bone interface produces partial reflection and transmission of the wave, preventing stresses from fully relaxing between reflections. This effect leads to a split wavefront, with faster propagation through the tooth region and slower propagation through the surrounding bone (**Fig. 6d**), resulting in increased reflection frequency and a more complex stress field. **Figure 6e** quantifies stress localization along two paths: through the tooth region and within the surrounding bone. Within the tooth pathway, the composite case exhibits the highest stresses, reaching peak compressive values of approximately 8 MPa. In contrast, stresses along the bone region remain comparable to the all-bone case (approx. 3.7 MPa compressive and 1.6 MPa tensile). These results indicate that the stacked tooth architecture localizes impact-induced stresses within the tooth material while limiting stress transmission into the supporting bone. Together, these findings show that material heterogeneity and geometric arrangement act to redistribute and confine impact energy, suggesting that stacked dentition provides a protective mechanism by concentrating stresses within expendable tooth material rather than the underlying skeletal structure.

**Figure 6.**
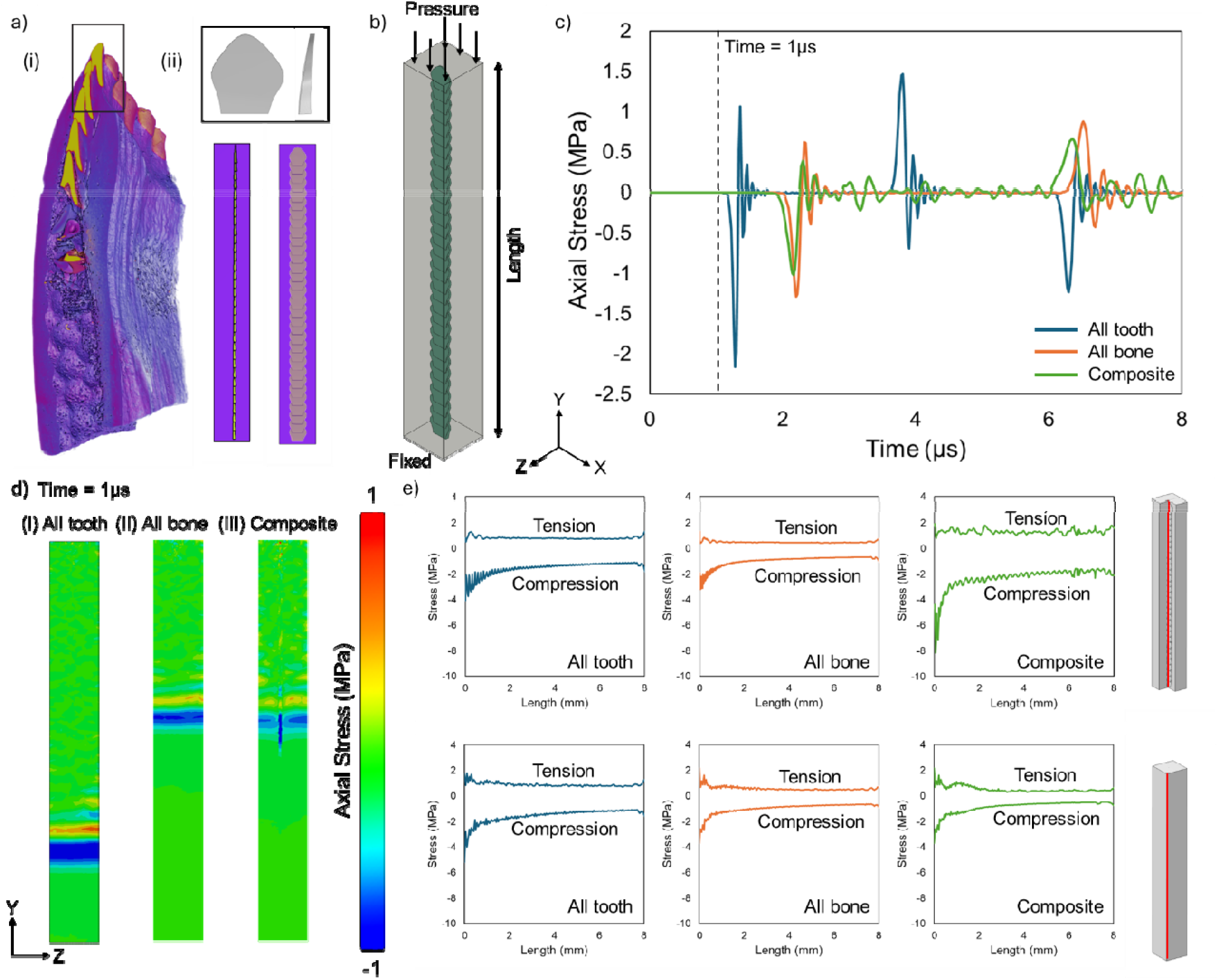
Shockwave propagation in a stacked tooth-bone parrotfish composite beak architecture. (a) Stacke tooth architecture in parrotfish beaks: (i) micro-computed tomography image showing stacked dentition, and (ii) simplified finite element model with a spade-shaped tooth embedded within bone. (b) Model geometry and boundary conditions for shockwave loading. A pressure impulse was applied at the top surface, the base was fixed, and lateral faces were constrained. (c) Average axial stress at the bottom face over 8 μs for three material configurations (all tooth, all bone and composite), showing differences in wave arrival and attenuation. (d) Axial stress contours for (i) all-tooth, (ii) all-bone and (iii) composite models. The composite case exhibits wavefront splitting due to differences in material properties between tooth and bone. (e) Maximum tensile and compressive axial stresses along the model length. Stresses within the tooth pathway (top) are elevated relative to those in the surrounding bone (bottom), indicating localization within the tooth region.

## 4 DISCUSSION

Parrotfish jaws represent a striking example of convergent evolution, characterized by fused beak-like dentition and stacked tooth architectures. These features have been linked to the ability to bite into coral, a mechanically hard biomaterial with stiffness spanning orders of magnitude (1-26 GPa^36^, 77 GPa^37^). The present results provide a quantitative biomechanical approach for understanding how these anatomical features influence mechanical performance under distinct loading conditions. The two loading regimes examined here, jaw opening and jawclosing, reveal fundamentally different mechanical behaviors. During jaw opening, deformation is governed by geometric scaling, with model size, hinge geometry and rotational inertia controlling displacement. In this regime, dentition morphology has minimal influence on global stiffness. In contrast, jawclosing imposes a constrained loading condition in which stress accumulation dominates the response. Under these conditions, fused dentition reduces the rate of stress development and promotes more uniform stress distribution, indicating that the advantage of the beak is specific to stabilized biting rather than unconstrained jaw motion.

### Foraging ecology influences beak performance

Foraging ecology plays an important role in the morphology and subsequent performance of feeding structures in vertebrates. In parrotfishes specifically, differences between scraping and excavating foraging ecologies may result in fine scale differences in morphology and performance among beaked species. In our study, convergently evolved excavating species *S. viride* and *C. ocellatus*, exhibit comparable stress responses despite significant phylogenetic separation^8^. This convergence in mechanical performance supports the interpretation that fused dentition represents a repeated functional solution to the demands of coral excavting, where resistance to localized stress accumulation is critical. Interestingly, the convergence in stress patterns closely mirrors the convergence in overall jaw shape between these species which has been noted in previous studies^12,18^. In contrast, the scraping *S. forsteni* shows divergent stress distributions relative to the other two beaked species and exhibits slightly higher peak stresses. Unlike excavating parrotfishes, scrapers do not remove substratum during their bites and feed at much faster rates than excavators. The differences in stress distributions between *S. forsteni* and the excavating parrotfishes may represent divergent functional demands as a result of different foraging ecologies. In the browsing parrotfishes, peak stresses were higher during controlled biting than the scraping and excavating species with stresses concentrated at the tips of their individual teeth. Unsurprisingly, this suggests that browsers likely feed in distinct ways on different food types relative to the other beaked species. In these browsing species, smaller flatter teeth are present toward the posterior margins of the jaws and previous studies have suggested that these teeth play an important role in shearing of pieces of macroalgae and seaweed during feeding while the more anterior teeth and plicidentine fangs may play a larger role in aggressive interactions between males^33^.

### The parrotfish beak as a shock absorber

The transient wave propagation simulations further show that tooth geometry governs not only stress magnitude but also the structure of propagating stress waves. Sharp-edged geometries produce elevated stress concentrations and pronounced tensile–compressive heterogeneity, whereas smooth and tapered profiles reduce peak stresses and disrupt coherent wave transmission. These results indicate that tooth shape modulates how impact energy enters the jaw. The consistent reduction in stress concentration for smooth profiles suggests that the absence of sharp corners is a key geometric feature, consistent with the prevalence of curved tooth morphologies across fish species^32^.

The stacked tooth architecture introduces an additional level of mechanical control through material heterogeneity. Differences in elastic properties between tooth and bone produce distinct wave speeds, resulting in wavefront splitting and partial reflection at the interface. This behavior localizes stresses within the tooth region while limiting transmission into the surrounding bone. Such localization is consistent with a protective mechanism in which expendable tooth material absorbs and redistributes impact energy, reducing mechanical demand on the underlying skeletal structure. Taken together, these results identify a coordinated biomechanical strategy in which geometry, morphology and material heterogeneity act across multiple length scale and time scales to control internal force transmission. Jaw opening is governed by geometric scaling, stabilized biting is governed by dentition morphology, and impact loading is regulated by tooth geometry and stacking architecture. This multiscale control suggests that convergent evolution in parrotfishes reflects optimization of stress redistribution and energy localization rather than simple structural increases in strength.

Several limitations should be considered. Material properties were assumed to be homogeneous and identical across species due to limited experimental data, including species-specific properties, which may obscure species-specific differences in composite structure. The models do not capture microstructural gradients and porosity at the tooth–bone interface, which likely influence wave propagation and stress transfer. Muscle forces were not explicitly modeled, and simplified boundary conditions may affect absolute stress magnitudes. In addition, the transient wave propagation models employ reduced-order geometries that do not capture full anatomical complexity or direct interaction with the coral substrate. However, the use of consistent assumptions across all cases enables robust comparison of relative trends across species and provides a framework for identifying how geometric and structural differences contribute to mechanical function. Future work should incorporate species-specific loading conditions, microstructural detail and explicit coral interaction to further refine these findings.

## 5 CONCLUSION

Parrotfishes are dynamic ecosystem engineers that have repeatedly evolved coalesced beaked morphologies to feed on hard substrates. Our results show that both the geometry and morphology of beaked dentitions and the individual teeth that comprise them can mitigate stresses during biting activities and absorb shockwaves by localizing wave transmissions to individual teeth which can be repeatedly replaced throughout the lifespans of these organisms. In contrast, non-beaked, browsing parrotfishes experienced more stress during simulated bites with stresses concentrated to individual teeth suggesting that their dental morphologies are substantially less optimized for durophagy. Taken together, our results suggest that parrotfish beaks are fine-tuned at multiple scales to reduce stress and absorb shock during biting and that strong selection for these attributes drove the convergent evolution of beaks within parrotfishes and perhaps the evolution of beaked dentition in other fishes more broadly.

## Acknowledgements

R.R.M acknowledges support from the Rice University Academy of Fellows. R.A. acknowledges the ASME Haythornthwaite Foundation Research Initiation Grants from the Applied Mechanics Division (AMD). K.M.E acknowledges Adam Summers for spirited discussions about parrotfish beaks, and Mark Westneat for additional parrotfish related inspiration and guidance. K.M.E. was supported by NSF DEB-2237278.

## FIGURES

**Supplementary Figure 1.**
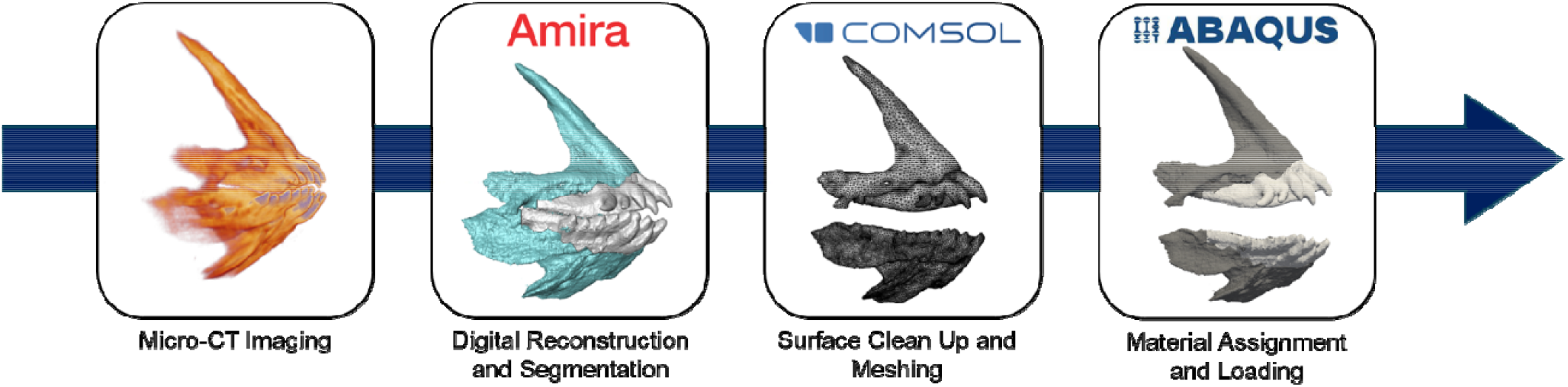
Digital reconstruction and model preparation pipeline. Step-by-step workflow for generation of finite element models, including imaging, segmentation, geometry processing and meshing.

**Supplementary Figure 1.**
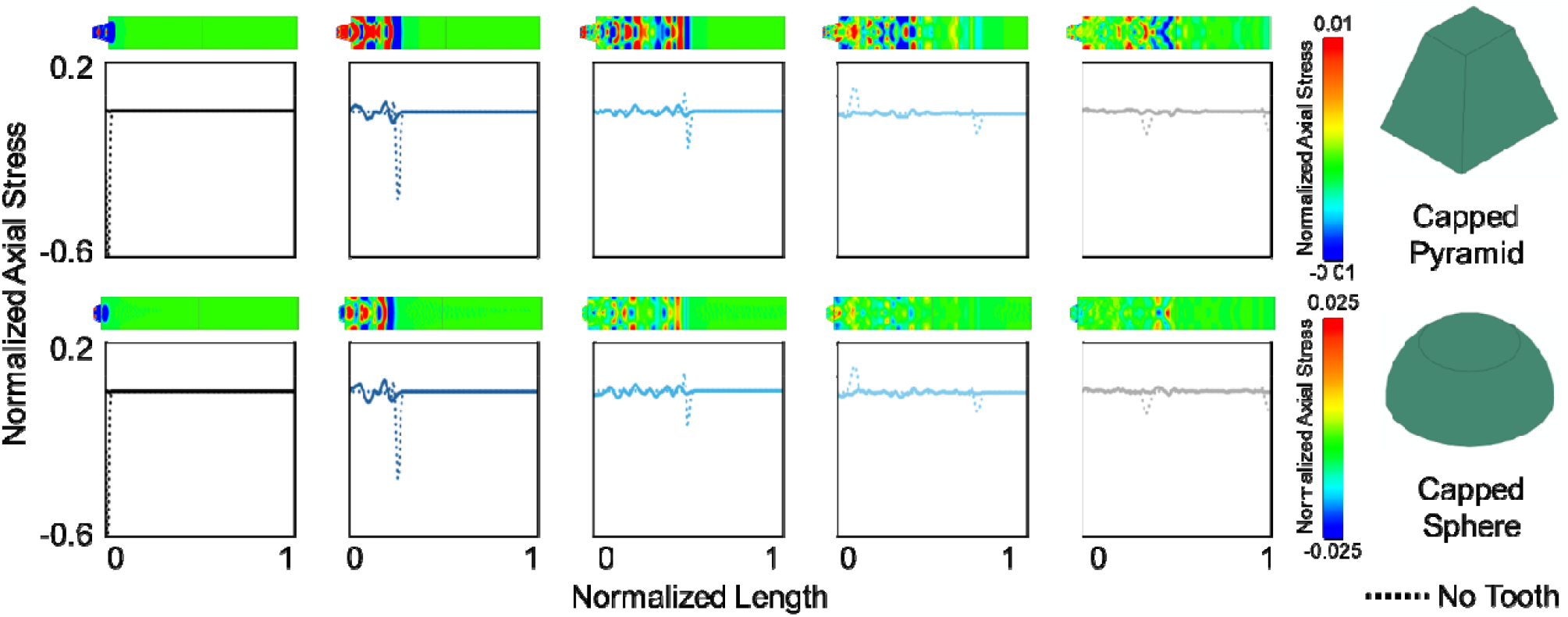
Normalized axial stress distributions for capped pyramid and capped spher geometries. Axial stress along the model length is normalized by the applied pressure to enable comparison across geometries. Solid lines show stress profiles for each geometry, with the no-tooth control shown as a dashed line. Columns correspond to tooth geometry and rows to successive time points; normalized axial stress contours are shown above each profile. Both geometries exhibit comparable normalized stress amplitudes. The capped pyrami shows greater spatial variation, with more pronounced alternating tensile–compressive regions than the cappe sphere.

**Supplementary Figure 2.**
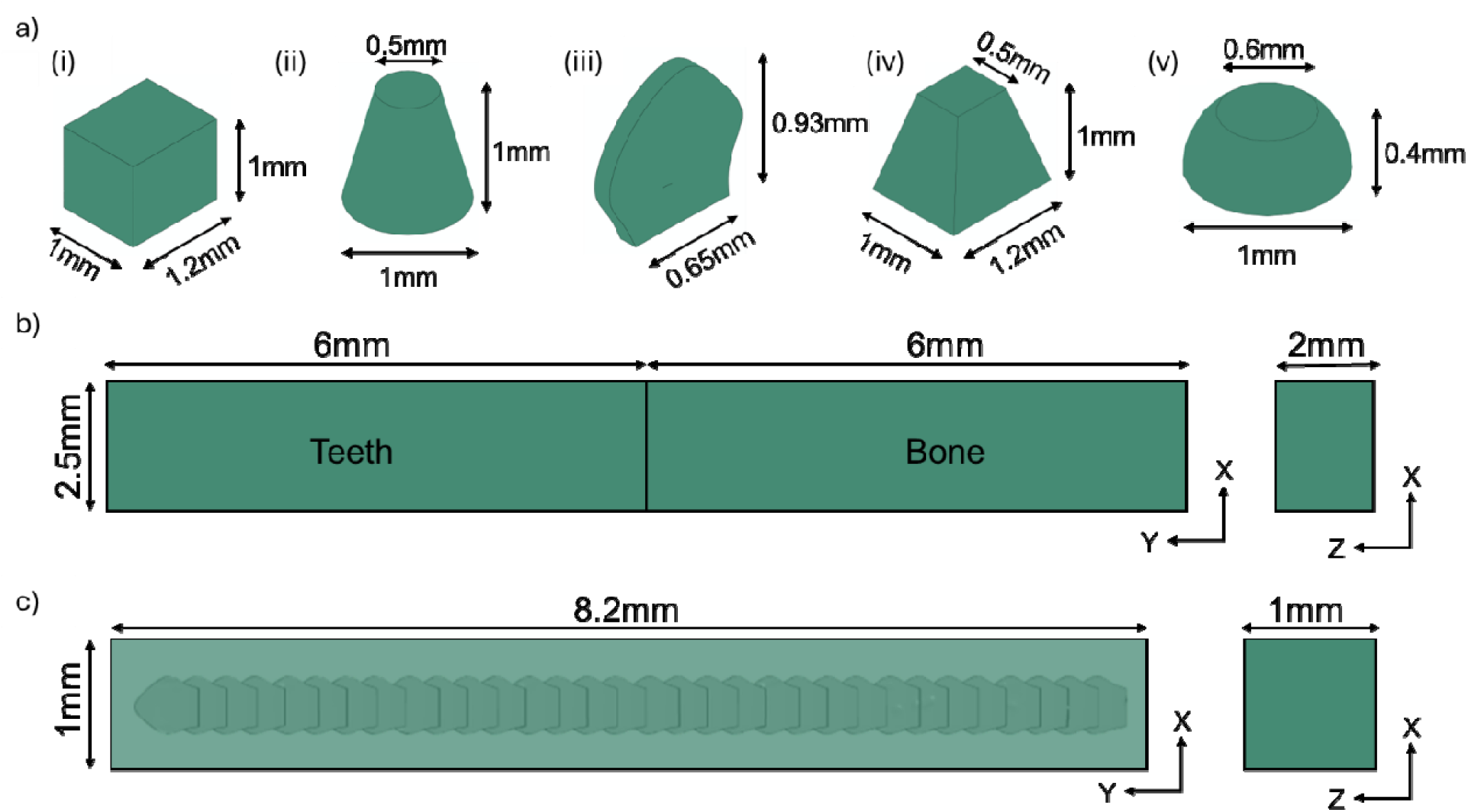
Geometries and dimensions of transient impact models. (a) Tooth profiles used in the impact analysis: (i) cuboid, (ii) frustum, (iii) spade, (iv) capped pyramid and (v) capped sphere. (b) Dimensions of the no-tooth control model. (c) Dimensions of the stacked tooth model.

**Supplementary Figure 3.**
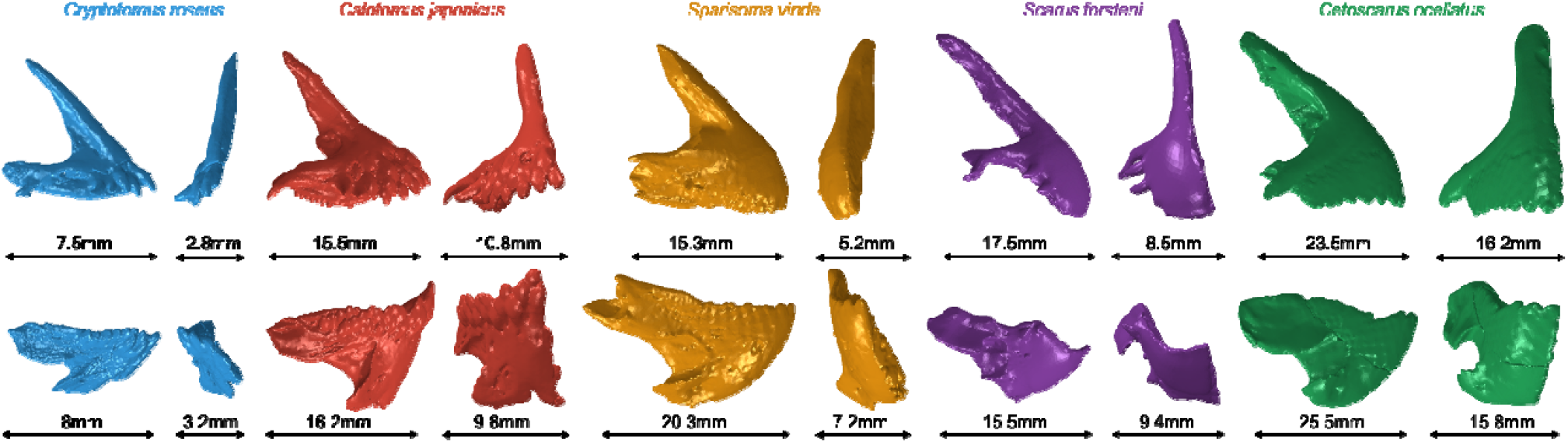
Jaw geometries and dimensions across parrotfish species. Side and front views of *Cryptotomus roseus, Cetoscarus ocellatus, Scarus forsteni, Sparisoma viride* and *Calotomus japonicus*, with characteristic dimensions indicated. *Cryptotomus roseus* exhibits the smallest jaw length (<10 mm), whereas *Cetoscarus ocellatus* exhibits the largest (>20 mm).

## Notes

### Competing Interest Statement

The authors have declared no competing interest.

## References

1. Ford, K. L., Peterson, R., Bernt, M. & Albert, J. S. Convergence is Only Skin Deep: Craniofacial Evolution in Electric Fishes from South America and Africa (Apteronotidae and Mormyridae). Integr Org Biol 4, obac022 (2022).

2. Winemiller, K. O. & Adite, A. Convergent evolution of weakly electric fishes from floodplain habitats in Africa and South America. Environmental Biology of Fishes 49, 175–186 (1997).

3. Keiler, J., Wirkner, C. S. & Richter, S. One hundred years of carcinization – the evolution of the crab-like habitus in Anomura (Arthropoda: Crustacea). Biol J Linn Soc 121, 200–222 (2017).

4. Wolfe, J. M., Luque, J. & Bracken-Grissom, H. D. How to become a crab: Phenotypic constraints on a recurring body plan. BioEssays 43, 2100020 (2021).

5. Alvarado-Cárdenas, L. O. et al. To converge or not to converge in environmental space: testing for similar environments between analogous succulent plants of North America and Africa. Ann Bot 111, 1125–1138 (2013).

6. Friedman, S. T., Price, S. A., Hoey, A. S. & Wainwright, P. C. Ecomorphological convergence in planktivorous surgeonfishes. j. evol. Biol. 29, 965–978 (2016).

7. Albertson, R. C. & Kocher, T. D. Genetic and developmental basis of cichlid trophic diversity. Heredity 97, 211–221 (2006).

8. Evans, K. M. et al. Beaks promote rapid morphological diversification along distinct evolutionary trajectories in labrid fishes (Eupercaria: Labridae). Evolution 77, 2000–2014 (2023).

9. Nicholson, G. M. & Clements, K. D. Micro-photoautotroph predation as a driver for trophic niche specialization in 12 syntopic Indo-Pacific parrotfish species. Biological Journal of the Linnean Society 139, 91–114 (2023).

10. Nicholson, G. M. & Clements, K. D. Ecomorphological divergence and trophic resource partitioning in 15 syntopic Indo-Pacific parrotfishes (Labridae: Scarini). Biological Journal of the Linnean Society 132, 590–611 (2021).

11. Bellwood, D. R. A Phylogenetic Study of the Parrotfishes Family Scaridae (Pisces: Labroidei): With a Revision of Genera. (Rodenprint, Sydney, 1994).

12. Wainwright, P. & Price, S. Innovation and Diversity of the Feeding Mechanism in Parrotfishes. in 26–41 (2018). doi:10.1201/9781315118079-2.

13. Adam, T. C., Kelley, M., Ruttenberg, B. I. & Burkepile, D. E. Resource partitioning along multiple niche axes drives functional diversity in parrotfishes on Caribbean coral reefs. Oecologia 179, 1173–1185 (2015).

14. Bonaldo, R. M., Welsh, J. Q. & Bellwood, D. R. Spatial and temporal variation in coral predation by parrotfishes on the GBR: evidence from an inshore reef. Coral Reefs 31, 263–272 (2012).

15. Ezzat, L. et al. Parrotfish predation drives distinct microbial communities in reef-building corals. Anim Microbiome 2, 5 (2020).

16. Bonaldo, R. M. & Bellwood, D. R. Parrotfish predation on massive Porites on the Great Barrier Reef. Coral Reefs 30, 259–269 (2011).

17. Clements, K. D., German, D. P., Piché, J., Tribollet, A. & Choat, J. H. Integrating ecological roles and trophic diversification on coral reefs: multiple lines of evidence identify parrotfishes as microphages. Biological Journal of the Linnean Society n/a,.

18. Streelman, J. T., Alfaro, M., Westneat, M. W., Bellwood, D. R. & Karl, S. A. Evolutionary History of the Parrotfishes: Biogeography, Ecomorphology, and Comparative Diversity. Evolution 56, 961–971 (2002).

19. Bellwood, D. R. & Choat, J. H. A functional analysis of grazing in parrotfishes (family Scaridae): the ecological implications. in Alternative life-history styles of fishes (ed. Bruton, M. N.) 189–214 (Springer Netherlands, Dordrecht, 1990). doi:10.1007/978-94-009-2065-1_11.

20. McCurry, M. R., Walmsley, C. W., Fitzgerald, E. M. G. & McHenry, C. R. The biomechanical consequences of longirostry in crocodilians and odontocetes. Journal of Biomechanics 56, 61–70 (2017).

21. Tseng, Z. J. Testing Adaptive Hypotheses of Convergence with Functional Landscapes: A Case Study of Bone-Cracking Hypercarnivores. PLoS One 8, e65305 (2013).

22. Amira Software | Biological Image Analysis Software - US. https://www.thermofisher.com/us/en/home/electron-microscopy/products/software-em-3d-vis/amira-software.html.

23. Marcus, M. A. et al. Parrotfish Teeth: Stiff Biominerals Whose Microstructure Makes Them Tough and Abrasion-Resistant To Bite Stony Corals. ACS Nano 11, 11856–11865 (2017).

24. Lai, Y.-S. et al. The Effect of Graft Strength on Knee Laxity and Graft In-Situ Forces after Posterior Cruciate Ligament Reconstruction. PLOS ONE 10, e0127293 (2015).

25. Hsieh, M.-T., Begley, M. & Valdevit, L. Architected implant designs for long bones: Advantages of minimal surface-based topologies. Materials & Design 207, 109838 (2021).

26. Rościszewska, M. & Zielinski, A. Structural and Material Determinants Influencing the Behavior of Porous Ti and Its Alloys Made by Additive Manufacturing Techniques for Biomedical Applications. Materials 14, 712 (2021).

27. Fluorapatite. https://www.mindat.org/min-1572.html.

28. Babaei, B. et al. The effect of dental restoration geometry and material properties on biomechanical behaviour of a treated molar tooth: A 3D finite element analysis. Journal of the Mechanical Behavior of Biomedical Materials 125, 104892 (2022).

29. Williams, K. L. & Price, S. A. Investigating Best Practices for Applying a Quantitative Tooth Complexity Metric to Fishes. Integr Comp Biol 65, 797–811 (2025).

30. Cohen, K. E., Weller, H. I. & Summers, A. P. Not your father’s homodonty—stress, tooth shape, and the functional homodont. J Anat 237, 837–848 (2020).

31. Hulsey, C. D. et al. Grand Challenges in Comparative Tooth Biology. Integr Comp Biol 60, 563–580 (2020).

32. Bellwood, D. R., Hoey, A. S., Bellwood, O. & Goatley, C. H. R. Evolution of long-toothed fishes and the changing nature of fish–benthos interactions on coral reefs. Nat Commun 5, 3144 (2014).

33. Viviani, J. et al. Plicidentine in the oral fangs of parrotfish (Scarinae, Labriformes). Journal of Anatomy 241, 601–615 (2022).

34. Melgarejo-Damián, M. P., González-Acosta, A. F., Cruz-Escalona, V. H. & Moncayo-Estrada, R. A comparison of feeding biomechanics between two parrotfish species from the Gulf of California. Zoomorphology 137, 165–176 (2018).

35. Gale, J. How strong is a parrotfish bite? The Institute for Environmental Research and Education https://iere.org/how-strong-is-a-parrotfish-bite/ (2025).

36. Wu, Z., Yu, H., Ma, H., Zhang, J. & Da, B. Physical and mechanical properties of coral aggregates in the South China Sea. Journal of Building Engineering 63, 105478 (2023).

37. Pasquini, L. et al. Isotropic microscale mechanical properties of coral skeletons. J R Soc Interface 12, 20150168 (2015).

